# Gut microbiota density influences host physiology and is shaped by host and microbial factors

**DOI:** 10.1101/277095

**Authors:** Eduardo J. Contijoch, Graham J. Britton, Chao Yang, Ilaria Mogno, Zhihua Li, Ruby Ng, Sean R. Llewellyn, Sheela Hira, Crystal Johnson, Keren M. Rabinowitz, Revital Barkan, Iris Dotan, Robert P. Hirten, Shih-Chen Fu, Yuying Luo, Nancy Yang, Tramy Luong, Philippe R. Labrias, Sergio A. Lira, Inga Peter, Ari Grinspan, Jose C. Clemente, Roman Kosoy, Seunghee Kim-Schulze, Xiaochen Qin, Anabella Castillo, Amanda Hurley, Ashish Atreja, Jason Rogers, Farah Fasihuddin, Merjona Saliaj, Amy Nolan, Pamela Reyes-Mercedes, Carina Rodriguez, Sarah Aly, Kenneth Santa-Cruz, Lauren A. Peters, Mayte Suárez-Fariñas, Ruiqi Huang, Ke Hao, Jun Zhu, Bin Zhang, Bojan Losic, Haritz Irizar, Won-Min Song, Antonio Di Narzo, Wenhui Wang, Benjamin L. Cohen, Christopher DiMaio, David Greenwald, Steven Itzkowitz, Aimee Lucas, James Marion, Elana Maser, Ryan Ungaro, Steven Naymagon, Joshua Novak, Brijen Shah, Thomas Ullman, Peter Rubin, James George, Peter Legnani, Shannon E Telesco, Joshua R. Friedman, Carrie Brodmerkel, Scott Plevy, Judy Cho, Jean-Frederic Colombel, Eric Schadt, Carmen Argmann, Marla Dubinsky, Andrew Kasarskis, Bruce Sands, Jeremiah J. Faith

## Abstract

To identify factors that regulate gut microbiota density and the impact of varied microbiota density on health, we assayed this fundamental ecosystem property in fecal samples across mammals, human disease, and therapeutic interventions. Physiologic features of the host (carrying capacity) and the fitness of the gut microbiota shape microbiota density. Therapeutic manipulation of microbiota density in mice altered host metabolic and immune homeostasis. In humans, gut microbiota density was reduced in Crohn’s disease, ulcerative colitis, and ileal pouch-anal anastomosis. The gut microbiota in recurrent *Clostridium difficile* infection had lower density and reduced fitness that were restored by fecal microbiota transplantation. Understanding the interplay between microbiota and disease in terms of microbiota density, host carrying capacity, and microbiota fitness provide new insights into microbiome structure and microbiome targeted therapeutics.

## Introduction

Population density is a fundamental parameter in understanding the health and function of any ecosystem, yet we know little about which host and microbial factors contribute to the density of organisms in the gut microbiota (i.e., gut microbiota density). The relationships uncovered between the gut microbiota and health over the past decade have largely focused on relative differences in community composition, estimated with culture-independent 16S rRNA gene (*1, 2*) or shotgun metagenomic sequencing (*3*). The microbiome’s influence on host physiology likely depends on the number – and not just the type – of bacteria interfacing with the host. Therefore, understanding factors driving gut microbiota density, as well as the impact of microbiota density on health, may advance the therapeutic potential of the microbiota.

Microbiota density has previously been measured with colony-forming units, DNA spike-ins (*4, 5*), qPCR (*6, 7*), flow cytometry (*8–10*), and microbial DNA quantification (microbial DNA per mass of sample) (*9, 11, 12*). Here, we use fecal microbial DNA content to estimate gut microbiota density, since it correlates with flow cytometry counts and colony forming units (CFU), and it can be easily incorporated into standard microbiome sequencing workflows by weighing the sample (*9*). We investigate host and microbial factors that contribute to microbiota density across a diverse set of mammalian microbiomes, study the impact of microbiota density on host adiposity and immune function in controlled mice models, and describe microbiota density changes in disease and the resolution of those alterations after therapy.

## Results

### The natural variation of gut microbiota density in mammals is driven by host and microbial factors

In macroecology, carrying capacity is the maximal density of organisms supported by an ecosystem and is broadly dictated by the resources (*e.g*. food, water, and habitat) in the environment. Whether or not the collection of species in an environment can reach the carrying capacity depends on their ability to efficiently utilize the available resources (i.e., the community’s fitness for the environment). To explore the contribution of host carrying capacity and gut microbiota fitness to microbiota density, we first collected fecal material from sixteen different mammalian species (Table S1) in order to sample a diverse range of host intestinal architectures and gut microbial community compositions. Using methods optimized to assay fecal microbiota density with greater throughput (see Methods and Figure S1), we observed significant differences in microbiota density across the mammalian samples (H = 69.0, p = 6.72 × 10^−9^; Kruskal-Wallis) with a 216-fold difference between the median of the most dense and least dense gut microbiota (Figure 1A). We found a positive correlation between microbiota density and phylogenetic relatedness of the host (R^2^ = 0.191; p = 0.0368; Pearson; Figure 1B), while there was no correlation between microbiota density and either fecal water content (*ρ* = −0.0418, p = 0.892, Spearman; Figure 1C) or host size (mass) (*ρ* = −0.364, p = 0.167, Spearman; Figure S2). Animals from order *Carnivora* (dog, ferret, lion, red panda, and tiger), with simple gut architectures adapted to carnivorous diets, had significantly reduced microbiota densities compared with the rest of the mammals studied (p = 6.14 × 10^−10^, Mann-Whitney, Figure 1D).

**Figure 1.**
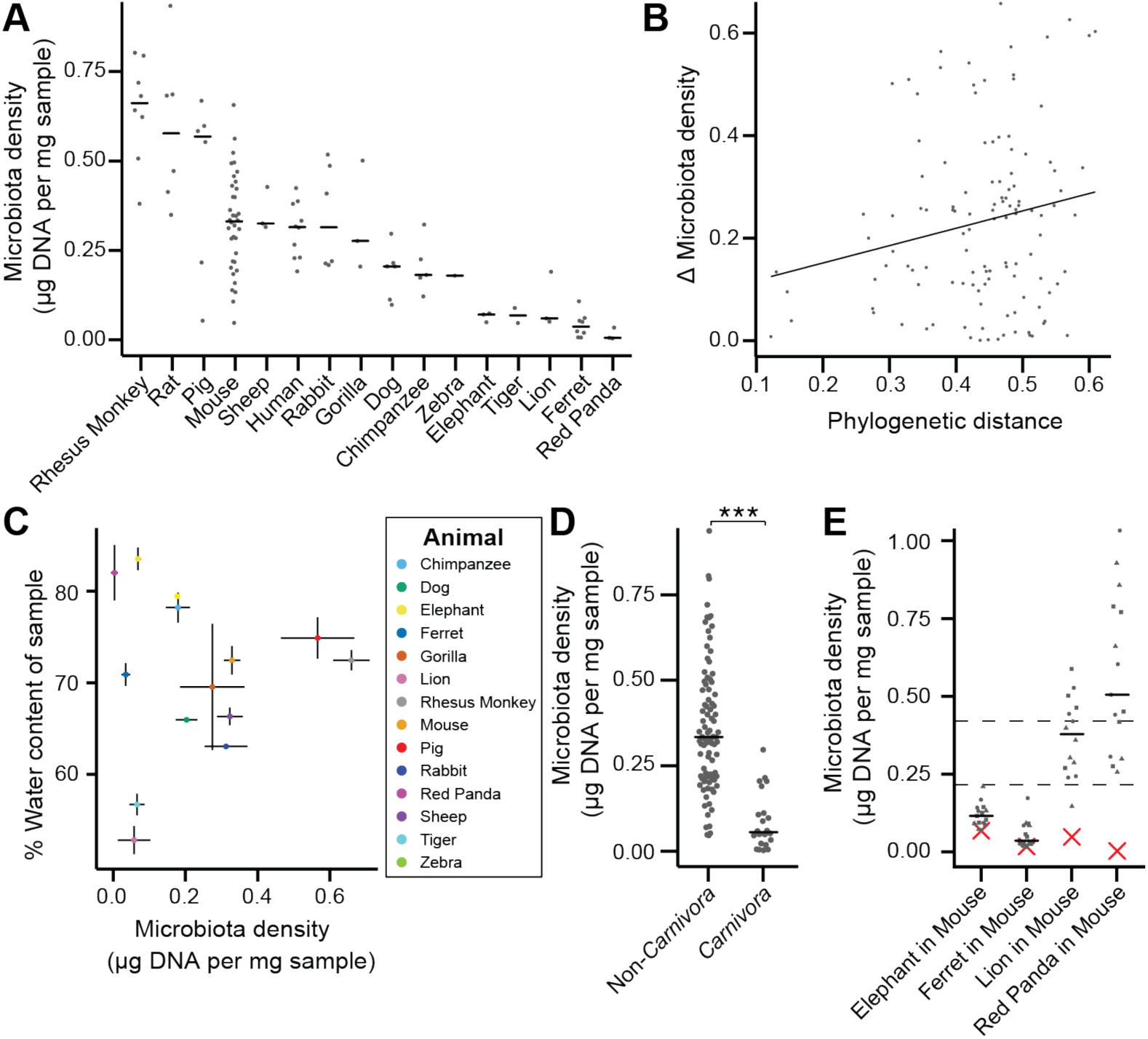
The natural variation in gut microbiota density across mammals is driven by host and microbial factors. (**A**) Fecal microbiota density varies across mammals. (**B**) Differences in microbiota density between animals was correlated with the degree of host phylogenetic relatedness as measured by mitochondrial 16S DNA sequence similarity. (**C**) Microbiota density and water content of fecal samples are not correlated. (**D**) Animals from the order *Carnivora* have a reduced microbiota density compared to mammals from other orders. (**E**) Different mammalian gut microbiotas transplanted into germ-free Swiss Webster mice (n = 3 per group) vary in their fitness to reach microbiota densities similar to mouse microbiotas. In **A** and **D-E**, points depict individual samples, and bars indicate median. In **C** points and lines indicate median values ± SEM. In **E** a red *X* indicates the microbiota density of the original mammalian sample, while dashed lines represent IQR of conventional Swiss Webster mice. ***p < 0.001.

To assay the relative contributions of the host (i.e., carrying capacity) and the microbiota (i.e., microbiota fitness) to microbiota density, we utilized germ-free mice with controlled host carrying capacity (*i.e.*, fixed diet, genetics, and environment) transplanted with the microbiotas of different mammals. Although there are clear caveats to assaying properties of the microbiota in a non-native host, several prior studies have demonstrated that germ-free microbiota transplantations from other mammals can recapitulate many aspects of the microbial community (*13–15*) and even host physiology (*16–19*) in the murine host. Importantly, these microbiota transplant experiments provide an experimental tool to estimate relative differences in fitness between microbiotas because each microbiota is transplanted into one or more replicate murine hosts with the same carrying capacity. In germ-free Swiss Webster mice colonized with four of the lowest density microbiotas in our initial screen (lion, elephant, ferret, and red panda), the lion and red panda microbiotas reached higher microbiota densities in the mouse than in the native host, suggesting their densities were limited by the carrying capacity of their host. The elephant and ferret microbiotas colonized mice at densities comparable to those in the native host and significantly less dense than a mouse microbiota (Figure 1E), suggesting their densities are limited by the fitness of each microbiota that cannot reach the mouse carrying capacity. Altogether, these mammalian microbiota samples and germ-free transfer experiments demonstrate that as in macroecology, microbiota density represents the combined influence of host carrying capacity and community fitness.

### Manipulation of colonic microbiota density alters host physiology

To broadly assess the impact of therapeutics on gut microbiota density, we provided SPF mice with one of 20 orally administered drugs, including antibiotics, anti-motility agents, and laxatives (Table S2). Only 9 of the 14 tested antibiotics significantly decreased gut microbiota density compared to untreated animals (p < 0.05 for each; Kruskal-Wallis rank sum test, followed by a Dunn’s test with Bonferroni correction). Amongst these 9 density-reducing antibiotics, there were substantial differences in each drug’s depleting capacity (Figure 2A). Of the laxatives, PEG 3350 reduced microbiota density (p = 2.22 × 10^−4^), while lactulose increased it (p = 0.0279). The anti-motility agent loperamide and the proton pump inhibitor omeprazole had no significant effect. Across the pharmacologics, we never observe high microbiota density with low alpha diversity, which drives a significant correlation between alpha diversity and microbiota density (*ρ* = 0.628, p < 0.0001, Spearman correlation; Fig. S3H). However, we commonly observe high alpha diversity with low microbiota density (e.g. animals given metronidazole; Fig. S3H), suggesting changes in microbiota density do not strictly correspond to changes in alpha diversity (see supplemental results and Fig. S3). As with our results in the mammals, we found no correlation between microbiota density and fecal water content across the tested pharmacologics (*ρ* = −0.338, p = 0.411, Spearman; Fig. S4).

**Figure 2.**
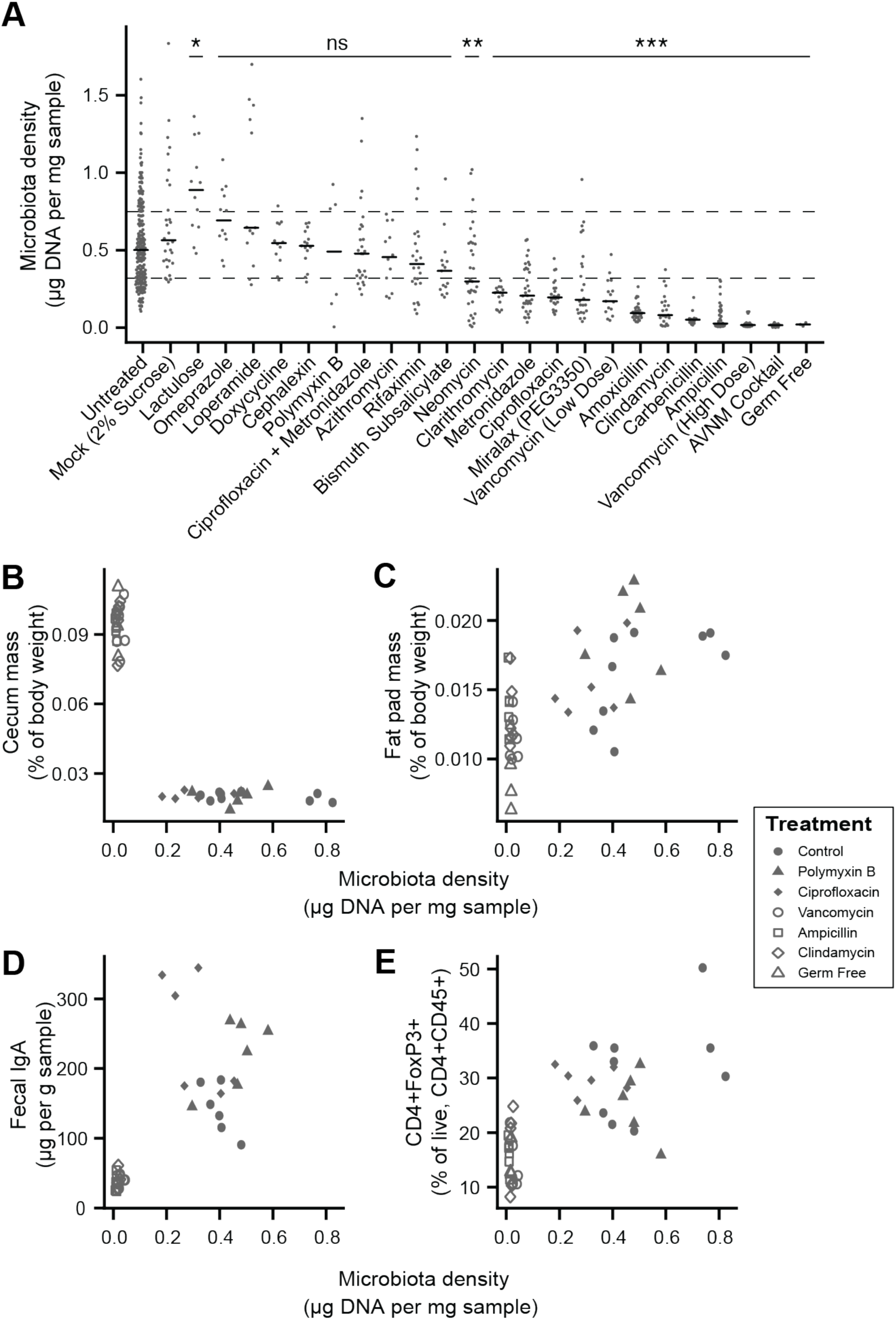
Manipulation of colonic microbiota density alters host physiology. (**A**) Pharmacologic interventions differentially alter microbiota density in SPF C57BL/6J mice. Samples from 3-12 (mean = 6) mice per group. (**B-E**) Antibiotic-induced changes in microbiota density significantly correlate with (**B**) host cecum size, (**C**) adiposity, (**D**) fecal IgA, and (**E**) colonic lamina propria FoxP3+ T regulatory cells. n = 6 mice per antibiotic group, 9 SPF antibiotic-free controls, and 6 germ-free controls. In **A**, dashed lines represent the IQR of untreated SPF C57BL/6J mice and AVNM = ampicillin, vancomycin, neomycin, metronidazole. Statistical tests performed for individual treatment conditions vs untreated using Kruskal-Wallis with Dunn’s post-test corrected for multiple comparisons with the Bonferonni correction. Bars indicate median. ns = not significant, *p < 0.05, **p < 0.01, and ***p < 0.001. In **B-E** points represent individual mice. Shapes indicate treatment group.

Comparing antibiotic-treated or germ-free mice with conventional mice has demonstrated the influence of the microbiota on a range of physiological measures (*15, 20–29*). To better understand the impact of microbiota density on host physiology, we selected five antibiotics (ampicillin, ciprofloxacin, clindamycin, polymyxin B, vancomycin) based on their varying ability to decrease microbiota density (Figure 2A). As expected, treating 4-week old SPF C57BL/6J mice with each antibiotic in their drinking water for four weeks (n = 6 mice per antibiotic, 9 SPF antibiotic-free controls, and 6 germ-free controls) led to a range of density reductions across the experimental groups (1.1 – 36.0 fold; Figure S4A). We found a significant negative correlation between cecum size and microbiota density (*ρ* = −0.729, p = 2.46 × 10^−7^, Spearman; Figures 2B and S4B). Epididymal fat pad mass, fecal IgA, and lamina propria FoxP3^+^CD4^+^ regulatory T cells were each positively correlated with microbiota density (*ρ*_fat_ = 0.587, p_fat_ = 6.11 × 10^−5^; *ρ*_IgA_ = 0.783, *ρ*_IgA_ = 3.35 × 10^−7^; *ρ*_Treg_ = 0.639, p_Treg_ = 5.31 × 10^−6^; Spearman; Figures 2C-2E and S4C-S4E). The strength of these associations is independent of the water content of the feces. Using group averages, the spearman correlations are the same for dry and wet microbiota density vs phenotypes (i.e., the rank order of density does not change when using dry weights). Furthermore, when estimating the relationships between microbiota density and host physiology with linear models we find that wet weight is a better predictor of changes in cecum size, epididymal fad pad mass, fecal IgA, and FoxP3^+^CD4^+^ regulatory T cells than dry weight.

### Microbiota density in inflammatory bowel disease (IBD)

To characterize the impact of host health status on gut microbiota density, we collected fecal samples from 70 healthy controls, 144 subjects with Crohn’s disease (CD), 109 subjects with ulcerative colitis (UC), and 19 subjects with UC that had undergone an ileal pouch-anal anastomosis (IPAA) procedure following total colectomy. Concordant with prior work using phylum-specific qPCR (*30*) and flow cytometry (CD-only; (*10*)), subjects with IBD had decreased microbiota density compared to healthy controls (p < 0.001 for each vs Healthy, Kruskal-Wallis rank sum test, followed by a Dunn’s test with Bonferroni correction; Figure 3A), even when excluding individuals receiving antibiotics (Figure S4A). Individuals with active CD or UC, as well as IPAA subjects had increased fecal water content compared to healthy individuals (p < 0.05 for each vs Healthy; Tukey’s HSD), while individuals with inactive CD or UC did not. Nonetheless, the decrease in microbiota density in IBD compared to healthy controls was consistent across individuals with active disease or inactive disease (p < 0.001 for each vs Healthy, Kruskal-Wallis rank sum test, followed by a Dunn’s test with Bonferroni correction; Figure 3B), demonstrating the microbiota density changes in IBD were not simply driven by the increased fecal water content that occurred with active inflammation.

**Figure 3.**
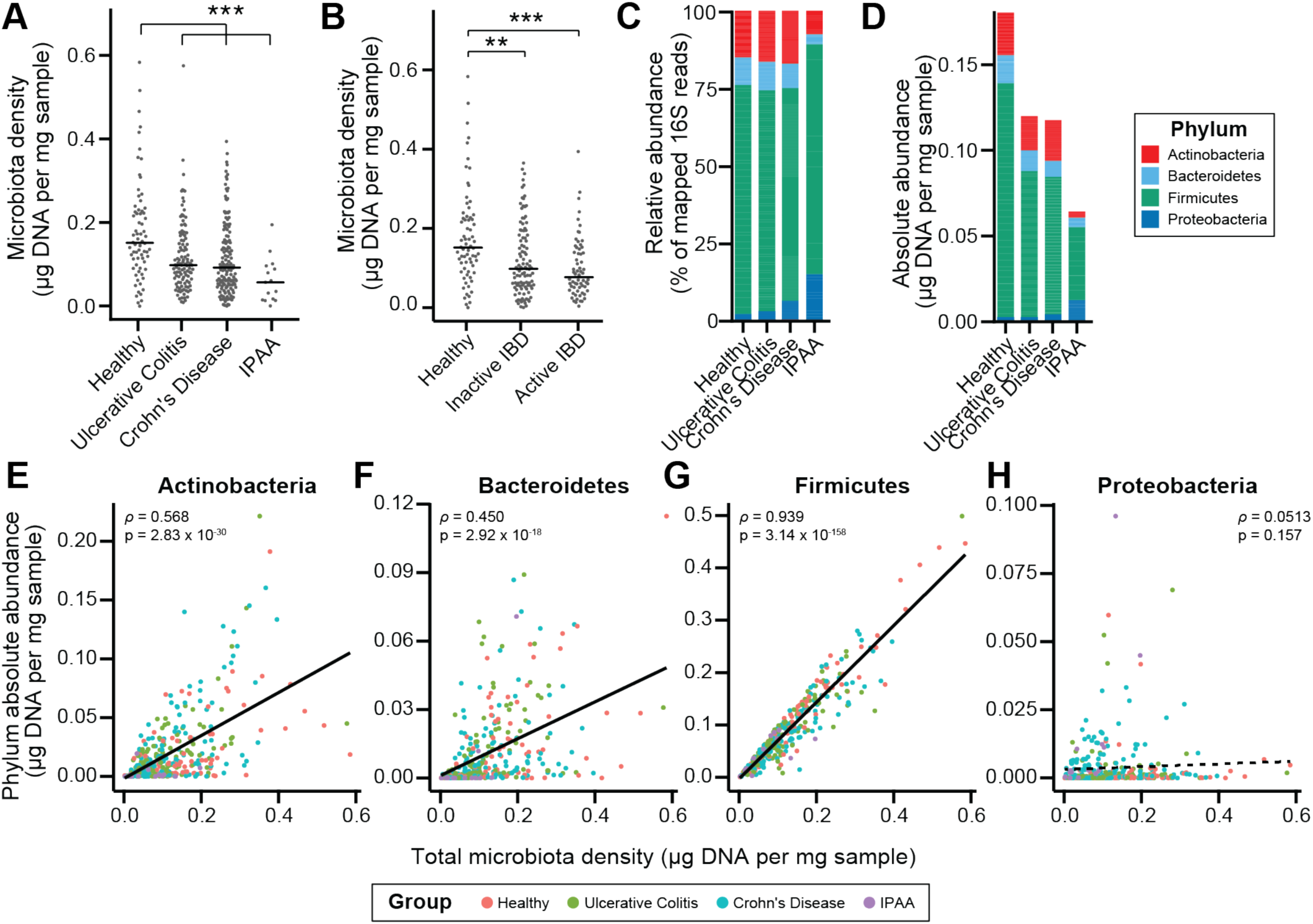
Microbiota density is altered in IBD. (**A**) Subjects with ulcerative colitis and Crohn’s disease, as well as subjects who have undergone ileal pouch-anal anastomosis (IPAA) have reduced microbiota density compared to healthy controls. (**B**) The reduction in microbiota density in IBD patients is independent of disease activity. (**C-D**) 16S rRNA gene sequencing reveals phylum-level changes in (**C**) relative and (**D**) absolute abundances of the microbiota in subjects with UC, CD, and IPAA compared to healthy controls. (**E-H**) The absolute abundance of all of the major phyla are strongly correlated with microbiota density, with the exception of Proteobacteria, whose abundance is largely constant. In **A-C**, bars indicate median, ** p < 0.01, and *** p < 0.001 (Kruskal-Wallis with Dunn’s post-test corrected for multiple comparisons with the Bonferonni correction). In **C**, each point represents the average microbiota density for an individual mouse before or after the initiation and development of colitis. In **E-H**, points represent individual subjects and colors indicate their health status.

To associate changes in microbiota composition with the altered microbiota density in individuals with IBD, we performed 16S rRNA gene amplicon sequencing of the fecal DNA (Figure 3C-3D). In line with previous studies (*30–33*), the IBD microbiome had a decreased alpha diversity compared to healthy subjects (p < 0.01 for all; Kruskal-Wallis rank sum test, followed by a Dunn’s test with Bonferroni correction; Figure S5). When we multiplied each taxa’s relative abundance by the microbiota density to calculate their absolute abundances, we found decreases in gut microbiota density were most significantly correlated with decreases in Firmicutes, while Proteobacteria were the only one of the four major phyla in the gut microbiota that were not correlated with microbiota density (Figures 3E-3H). These results from measuring the absolute microbiota differ from common observations of relative increases in Proteobacteria associated with IBD (*30, 31*). More accurately, it appears that Proteobacteria are able to sustain a constant density in the in IBD as the other phyla decrease in absolute terms.

### Fecal microbiota transplants restore microbiota density and microbiota fitness

Given the large difference in the microbiota between healthy individuals and those with recurrent *Clostridium difficile* infection (rCDI) (Figure S6; (*34, 35*)), we hypothesized that on a mechanistic level, FMT bolsters colonization resistance by improving gut microbiota fitness. In fecal samples from FMT donors and their rCDI FMT recipients prior to and after FMT, we observed that the rCDI gut microbiota has a significantly lower microbiota density than the donor microbiota, and that FMT increased microbiota density (p < 0.05 for all comparisons, Kruskal-Wallis rank sum test, followed by a Dunn’s test with Bonferroni correction; Figure 4A). We did not observe any differences in fecal water content between the donors and recipients before or after FMT (p > 0.2 for all comparisons, Tukey’s HSD). In addition, we found that rCDI FMT recipients had both a relative and absolute increase in Proteobacteria that was significantly reduced by FMT (Figures 4B, 4C, and S6C-S6F). These data suggest that FMT restores higher densities of Bacteroidetes, Firmicutes, and Actinobacteria to more fully realize the host’s carrying capacity. However, several factors may confound this conclusion as individuals with rCDI are often on antibiotics prior to FMT.

**Figure 4.**
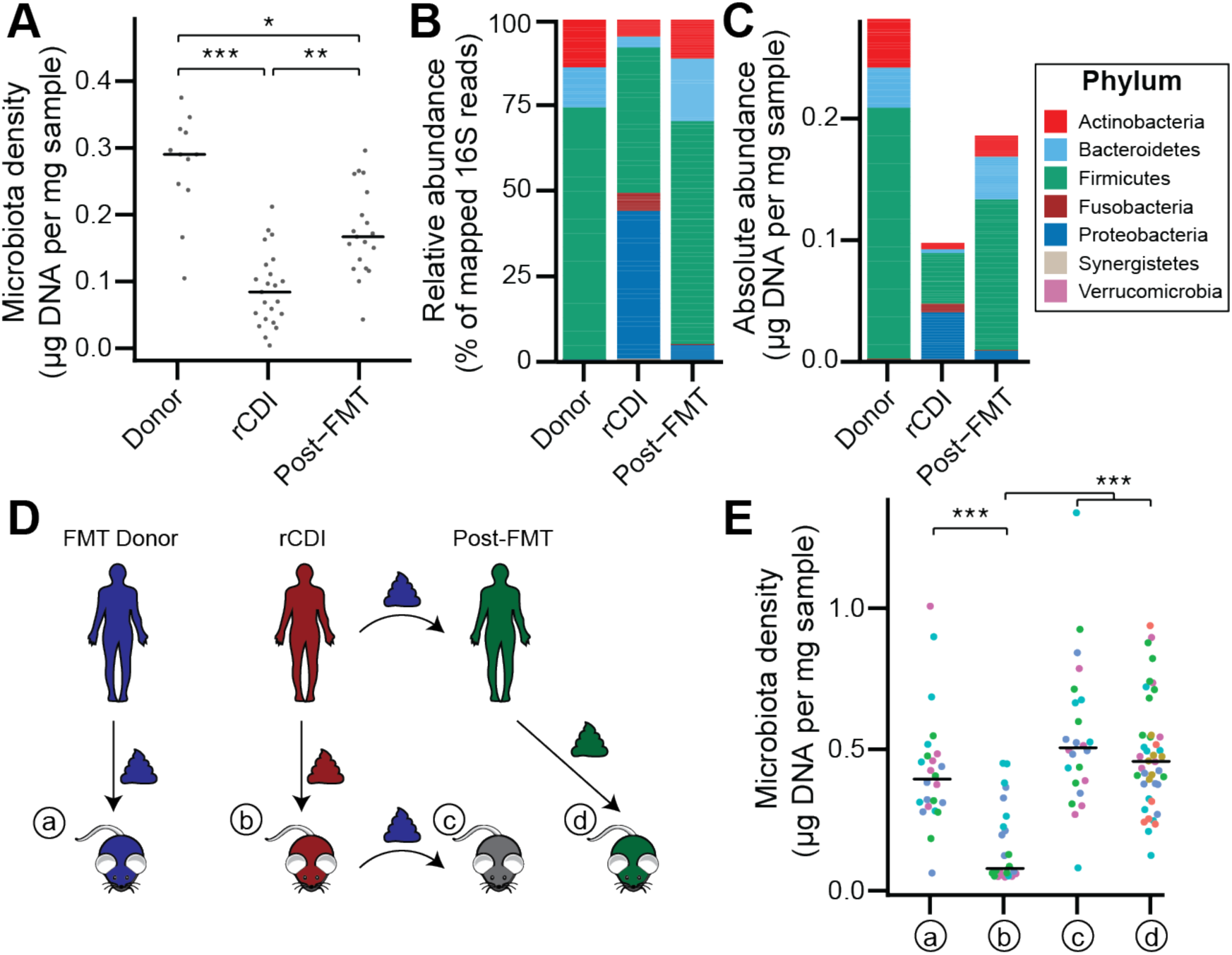
The rCDI microbiota has a fitness defect that is therapeutically treatable by FMT. (**A**) rCDI subjects have reduced microbiota densities that are significantly increased upon FMT with donor microbiotas. (**B** and **C**) Following FMT, the composition of the microbiota of individuals with rCDI is restored to more closely resemble that of healthy donors in both (**B**) relative and (**C**) absolute terms. (**D**) Germ-free mice were colonized with the microbiota from FMT Donors (a), or individuals with rCDI that underwent FMT (b). These mice then received the microbiota from the FMT donor corresponding to the clinical FMT (c), which could be compared to germ-free mice colonized with the Post-FMT sample from the individual who received the FMT (d). (**E**) Microbiota density in mice from the experimental scheme described in (**D**) showed decrease in microbiota fitness prior to FMT and an increase in microbiota density following FMT demonstrating the restoration of community fitness. In **A** and **E** points represent individual samples, bars indicate median, *p < 0.05, **p < 0.01, and ***p < 0.001 (Kruskal-Wallis with Dunn’s post-test corrected for multiple comparisons with the Bonferonni correction). In **E**, colors represent each one of five different FMT donor-recipient pairs.

To separate the host physiologic and pharmacologic factors that might impact our understanding of community fitness in rCDI, we utilized a gnotobiotic murine model of FMT (Figure 4D) where germ-free mice were initially colonized with the fecal material of individuals with rCDI for 3 weeks prior to a single transplant of fecal material via oral gavage from a second human donor – the same healthy FMT donor used for the transplant clinically. As a control, we colonized germ-free mice with the FMT donor microbiota alone (Figure 4D). The microbiota density of mice colonized with the healthy samples (a) was greater than that of mice colonized with rCDI samples (b) (p < 0.001, Kruskal-Wallis rank sum test, followed by a Dunn’s test with Bonferroni correction; Figure 5E), suggesting that rCDI individuals have a reduced microbiota fitness compared to healthy donors. Following the introduction of the healthy donor microbiota to the mice colonized with the rCDI microbiota (c), we observed increased microbiota density in these mice (p < 0.001, Kruskal-Wallis rank sum test, followed by a Dunn’s test with Bonferroni correction; Figure 4E), implying a restoration of microbiota fitness. Furthermore, when we colonize germ-free mice with the microbiota of the individuals with rCDI 6-12 months after they received an FMT (d), we find that their microbiota fitness had been restored, just as in our mouse FMT model (p < 0.001, Kruskal-Wallis rank sum test, followed by a Dunn’s test with Bonferroni correction; Figure 4E). These findings in the mice model recapitulate the data in our human cohort of FMT recipients and suggest that FMT successfully treats the fitness defect of the rCDI community.

## Discussion

The DNA-based microbiota density estimation method employed here and in previous studies (*9, 11, 12*) has the advantage that it can be incorporated into existing 16S rRNA and metagenomic workflows by simply weighing the input sample and ensuring the input mass of fecal material is within the linear range of the DNA extraction protocol. Incorporating microbiota density into standard culture independent microbiome workflows would greatly broaden our understanding of factors that drive one of the most fundamental properties of any ecosystem – its population density – and it would allow the broader study of absolute taxon abundances. Recent work has demonstrated that the amount of live/dead bacteria can vary between fecal samples (*36–38*), which would not be captured by a DNA-based density metric. However, in practice we found that the influence of any variation from live/dead bacteria was sufficiently low that it did not influence the major conclusions of this study; we observed a very significant correlation between the viability-based CFU density measurement and the DNA-based one and all of the major relationships observed in this study were consistent across both approaches (i.e., variation across mammals, IBD and IPAA lower density than healthy, rCDI lower density than FMT donor or rCDI post-transplant; Figure S1D).

Previous work has demonstrated that changes in fecal water are associated but not necessarily causally influencing differences in microbiota composition across the human population (*39*, *40*). While both fecal water content and microbiota density vary across mammals and can be altered by pharmacologics (Fig. 2, S4F), dietary components (Fig. S7; (*12*)), and host disease status, we find microbiota density is consistently not correlated with water content. In the context of altering host physiology through antibiotic manipulation of microbiota density, the best predictor of the impact of changes of microbiota density on host physiology was when density was calculated with stool wet weight, suggesting both wet and dry components of stool are important diluents in determining microbiota density and its impact on the host.

Differences in microbiota density can be influenced by both the host’s carrying capacity and the fitness of the microbiota to reach the carrying capacity of a given host. We found the density of gut microbes varies across mammals and is more similar in more phylogenetically related species. Across mammals, gut architecture appears to be a major driver of density, as the lowest densities were observed in order *Carnivora*, whose short, simple intestines have a lower carrying capacity and are maladapted for microbial fermentation at high densities. The low microbiota density of the red panda, a member of *Carnivora* with a herbivorous diet, further supports intestinal architecture as a major determinant of host carrying capacity and thus a driver of microbiota density. Finally, the significantly reduced microbiota density in humans with IPAA uniquely demonstrates that changing gut architecture within a species (in this case by surgery to treat ulcerative colitis) is equally capable of influencing host carrying capacity.

Within a murine host with controlled carrying capacity (i.e., fixed diet, genetics, housing, etc.), we found microbiota density can be altered with pharmacologics, with downstream consequences to host adiposity and immune function. Different antibiotics were highly varied in their ability to impact microbiota density, which could explain the mixed efficacy of antibiotics in microbiota-targeted clinical trials for complex disease and varied responses to antibiotics in animal models. Identifying more effective microbiota depleting cocktails would improve the design of such studies, while measuring microbiota density in trials with antibiotics could better stratify clinical response. Previous studies have observed that microbiota density can be manipulated by dietary changes (*12, 41*). Furthermore, we found that altering microbiota density with either diet or antibiotics could modify colitis severity (*12*). Understanding the long-term effect of high or low microbiota density on health could help refine the use of diet and the microbiota in disease treatment and prevention.

Finally, we observed that microbiota density is reduced in both IBD and rCDI. Microbiota density reductions, from a lack of fitness in the microbial community, were “druggable” by FMT. The ability of FMT to increase microbiota density through improved community fitness provides mechanistic insights into FMT for rCDI and a novel biomarker to track its success. This result also highlights that routine monitoring to identify individuals with microbiota fitness deficiencies combined with prophylactic microbial therapeutics targeted might form a therapeutic strategy to boost colonization resistance to treat or prevent disease.

## Acknowledgements

We are grateful to C. Fermin, E. Vazquez, and G. Escano in the Mount Sinai Immunology Institute Gnotobiotic facility for their help with gnotobiotic animal husbandry. D. Present and S. Petrunio provided helpful suggestions during the course of this work. Next generation sequencing was performed at NYU School of Medicine by the Genome Technology Center partially supported by the Cancer Center Support Grant, P30CA016087. Human microbiome processing was performed in part by the Human Immune Monitoring Center at the Icahn School of Medicine at Mount Sinai. This work was supported in part by the staff and resources of Scientific Computing and of the Flow Cytometry Core at the Icahn School of Medicine at Mount Sinai. Funding: This work was supported by grants from the Leona M. and Harry B. Helmsley Charitable Trust (I.D.), the NIH (NIGMS GM108505 for J.J.F. and NIDDK DK112679 for E.J.C.), Janssen Research & Development LLC, and SUCCESS. Raw sequencing files (fastq) for all 16S sequencing samples (antibiotic-treated mice, IBD cohort, and rCDI FMT cohort) are stored in the public Sequence Read Archive (SRA) under project number PRJNA413199. Flow cytometry data has been uploaded to Mendeley Data (http://dx.doi.org/10.17632/cjvfrbyxhj.1).

## Supplemental Methods

### Mammalian samples

Fecal samples from the mammals used in this study were collected either from laboratory animals housed and maintained at the Icahn School of Medicine at Mount Sinai (New York, NY), or from animals at the Zoo Knoxville (Knoxville, TN). Approximate animal masses were curated from the literature (*42–47*).

### Mice

Specific pathogen free (SPF) mice were purchased from Jackson Labs (C57BL/6J mice) or Taconic (Swiss Webster mice). Germ-free (GF) WT and Rag1^−/−^ C57BL/6J, and Swiss Webster mice were housed in standard, commercially available flexible film isolators. To generate gnotobiotic mice from human or mammalian fecal samples, GF mice were gavaged with 200 *μ*L of clarified stool from the source. Four week old male mice were used for the antibiotic treatment phenotyping experiments (Figure 3). All other experiments used both male and female mice between 4 and 6 weeks old. Swiss Webster mice were used to perform gnotobiotic experiments. Experiments and animal care were performed in accordance with the policies of the Icahn School of Medicine Institutional Animal Care and Use Committee (IACUC).

### Human subjects

To study the microbiota of individuals with IBD, we collected fecal samples from 70 healthy controls (42 female, 28 male), with an average age of 55.1 (range: 23-73), 109 individuals with ulcerative colitis (67 female, 42 male), with an average age of 52.8 (range: 22-80), and 144 individuals with Crohn’s Disease (72 female, 72 male), with an average age of 41.7 (range: 22-79). The study was approved by the Institutional Review Board (IRB) at Mount Sinai. For subjects with ulcerative colitis we defined disease activity using the Mayo Endoscopic Subscore (Mayo). Individuals with a Mayo = 3 were categorized as having active disease, and individuals with a Mayo = 0 were categorized as having inactive disease. For individuals with Crohn’s disease, active disease was defined as a Simple Endoscopic Score for Crohn Disease (SES-CD) ≥ 5, and inactive disease as SES-CD = 0. The remaining samples were excluded from these analyses. Stool samples were also collected from individuals with ulcerative colitis that had undergone an ileal pouch-anal anastamosis procedure following total colectomy (3 female, 12 male), with an average age of 42.93 (range: 19-68). These samples were collected from individuals in accordance with the IRB at the Tel Aviv Sourasky Medical Center. All individuals signed an informed consent. For the analysis of the change in the microbiota in recurrent *Clostridium difficile* infection following fecal microbiota transplantation, we collected samples from 11 healthy donors (8 female, 3 male; average age: 47.9, range: 25-75), 12 recipients who also had IBD (8 female, 4 male; average age: 55.3, range: 32-78), and 11 recipients who did not have IBD (9 female, 3 male; average age: 62, range: 36-87), as described in (*48*). The study was approved by the Mount Sinai IRB.

### Fecal sample collection and pre-processing

To quantify the mass of each fecal sample or fecal sample aliquot, we pre-weighed tubes prior to sample collection and post-weighed the tubes after adding the fecal material. For mouse samples, fresh fecal samples were collected directly into the collection tubes and stored at −80°C. For all other mammalian species with larger fecal sample sizes, samples were aliquoted on dry ice or liquid nitrogen and stored at −80°C. Sample aliquot sizes were targeted in the linear range of the fecal DNA extraction protocol (approx. 50 mg in mice and <200 mg in humans) to enable quantitative yields of DNA from the fecal material.

### Phenol:chloroform DNA extraction

Fecal samples processed with the phenol:chloroform DNA extraction method were collected into 2.0 mL collection tubes (Axygen, SCT-200-SS-C-S). Similar to previous studies (*9*), samples were suspended in a solution containing 282 *μ*L of extraction buffer (20 mM Tris (pH 8.0), 200 mM NaCl, 2mM EDTA), 200 *μL* 20% SDS, 550 *μ*L phenol:chloroform:isoamyl alcohol (25:24:1, pH 7.9), and 400 *μ*L of 0.1 mm diameter zirconia/silica beads (BioSpec, 11079101z). Samples were then lysed by mechanical disruption with a Mini-Beadbeater-96 (BioSpec, 1001) for 5 minutes at room temperature. Samples were centrifuged at 4000rpm for 5 minutes to separate aqueous and organic phases. The aqueous phase was collected and mixed with 650 *μ*L of PM Buffer (Qiagen, 19083). DNA extracts were then purified using a Qiagen PCR Purification kit (Qiagen, 28181), and eluted into 100 *μ*L of EB buffer. Purified DNA was quantified using the Broad Range or High Sensitivity Quant-IT dsDNA Assay kit (Thermo Fisher, Q32853 and Q33130) in combination with a BioTek Synergy HTX Multi-Mode Reader.

### DNase Inactivation Buffer DNA extraction

Phenol:chloroform based DNA extraction with bead beating is an effective method to isolate microbial DNA from feces. However, automation of phenol:chloroform requires liquid handling robotics in an environment compatible with this hazardous chemical mixture. In addition, the variable volume of the aqueous phase produced with this method presents an obstacle for its automation. We therefore tested the DIB bead beating extraction protocol as an alternative, since by eliminating the hazardous chemicals the protocol is compatible with more high-throughput liquid handling robotics platforms. Samples processed with the DNase Inactivation Buffer (DIB) DNA extraction method were collected into 1.0 mL tubes (Thermo Fisher, 3740). Samples were suspended in a solution containing 700 *μ*L of DIB (0.5% SDS, 0.5 mM EDTA, 20 mM Tris (pH 8.0)) and 200 *μ*L of 0.1 mm diameter zirconia/silica beads. Samples were then lysed by mechanical disruption and centrifuged as above. Since there is no phase separation with this method, it is straightforward to subsample the supernatant to improve the dynamic range of DNA quantification by avoiding saturating the column with DNA quantities above the binding capacity. 50-200 *μ*L of the supernatant was transferred into new collection tubes. Depending on the volume collected, an additional volume of DIB was added in order to reach a total volume of 200 *μ*L. Next, this DIB lysate was combined with 600 *μ*L of PM Buffer, purified with a Qiagen PCR Purification kit, and eluted into 100 *μ*L of EB buffer. Purified DNA was quantified using the Broad Range or High Sensitivity Quant-IT dsDNA Assay kit in combination with a BioTek Synergy HTX Multi-Mode Reader.

### 16S rRNA sequencing

DNA templates were normalized to 2 ng/*μ*L, and the V4 variable region of the 16S rRNA gene was amplified by PCR using indexed primers as previously described (*49*). The uniquely indexed 16S rRNA V4 amplicons were pooled and purified with AmpureXP beads (Beckman Coulter) with a ratio of 1:1 beads to PCR reaction. Correct amplicon size and the absence of primer dimers were verified by gel electrophoresis. The pooled samples were sequenced with an Illumina MiSeq (paired-end 250bp).

### Fecal sample water content

Samples were collected into pre-weighed 2.0 mL collection tubes (Axygen, SCT-200-SS-C-S). After collecting a fecal sample, sample mass was determined by post-weighing the tube. To measure the water content of a sample, tubes were placed at 105°C for 24 hours, and weighed again (*50*). The water content of a sample was calculated as the difference in final and initial mass of the sample, divided by the initial mass.

### Pharmacologic treatment of mice

Antibiotics (and other compounds) were provided *ad libitum* to mice in their drinking water, when possible. All of the pharmacologics were prepared into a 2% sucrose solution (which also served as the control treatment) and sterilized with a 0.22 *μ*m filter. Compounds that were not readily water-soluble were administered to mice via oral gavage of 200 *μ*L once per day, as indicated in Table S2. Unless identified otherwise, antibiotic and pharmacologic concentrations were calculated using a maximal clinical dose (taken from the online clinical resource UpToDate.com) or from previous studies (*20, 51–54*), assuming a 20 g mouse that drinks 3 mL water per day.

### Measurement of fecal immunoglobulin A

Fecal pellets were collected and massed. To each fecal pellet, 1 mL of sterile PBS was added per 100 mg feces. Each sample was homogenized without beads in a Mini-Beadbeater-96 for 3 min (BioSpec, 1001) followed by vortexing for 3 min. Samples were centrifuged at 9000g for 10 min at 4°C and supernatants were collected. Immunoglobulin A was measured by ELISA. Plates were coated with a working concentration of 1 ng/*μ*L of goat anti-mouse IgA-UNLB (SouthernBiotech Cat# 1040-01, RRID:AB_2314669), and then blocked with 1% BSA in PBS overnight at 4°C. Wells were washed with washing buffer (0.1% Tween-20 in PBS) 3 times. Then, fecal supernatant was diluted in dilution buffer (0.1% Tween-20, 1% BSA in PBS), added to each well, and incubated overnight at 4°C. The wells were washed again with washing buffer 5 times, and incubated for 2 hours at room temperature with a 1/2000 dilution of goat anti-mouse IgA-HRP (Sigma-Aldrich Cat# A4789, RRID:AB_258201) in dilution buffer. Following the incubation, the wells were washed 5 times with PBS/Tween-20. Next, TMB substrate was added to wells for 1 minute (KBL, 50-76-02 and 50-65-02), and the reaction was quenched using 1M H_2_SO_4_. Absorbance at 450 nm was measured using a BioTek Synergy HTX Multi-Mode Reader. Samples were quantified against a standard curve from 1000 ng/mL to 0.5 ng/mL.

### CFU Assay

We performed colony forming unit assays to obtain a culture-dependent measurement of microbiota density that also incorporates viability, as only live microbes will form colonies in this assay. Fecal samples were stored at −80°C after sampling. Prior to plating larger samples were pulverized under liquid nitrogen. Approximately 500mg of fecal sample was homogenized in 12 ml of rich broth and filtered with a 100 uM filter to remove particulate matter (*16*). Serial dilutions of this clarified fecal slurry were plated on chocolate agar and grown in an anaerobic chamber at 37°C for 72hr, whereupon colonies were manually quantified and normalized to CFU/g feces.

### Colonic lamina propria immune populations

Colonic lamina propria immune cell populations were measured as previously described (*16*). Briefly, colonic tissue was dissected and placed into RPMI medium at 4°C. Tissues were then transferred into HBSS and vortexed briefly, before being transferred into dissociation buffer (10% FBS, 5 mM EDTA, 15 mM HEPES in HBSS) and shaken for 30 minutes at 110 rpm at 37°C. Tissues were washed in HBSS before digestion in HBSS containing 2% FBS, 0.5 mg/mL Collagenase VIII (Sigma C2139) and 0.5 mg/mL DNase 1 (Sigma DN25) for 30 minutes at 110 rpm at 37°C. Digested tissue was then passed through a 100 *μ*m filter into cold RPMI medium. Samples were then centrifuged at 1500 rpm, 4°C for 5 minutes. The supernatant was removed and cells were washed once more in PBS before staining for flow cytometry. No enrichment of mononuclear cells by density centrifugation was performed. Cells were initially blocked with Fc Block (BioLegend Cat# 101320, RRID:AB_1574975) and subsequently stained for: viability (BioLegend Cat# 423101) and immunolabelled for expression of CD4 (1:200, BioLegend Cat# 100411, RRID:AB_312696) and CD45 (1:100, BioLegend Cat# 103115, RRID:AB_312980), and FoxP3 (1:100, Thermo Fisher Scientific Cat# 12-5773-82, RRID:AB_465936). Surface markers were stained before fixation and intracellular markers were stained after fixation with the FoxP3 Fixation/Permeabilization Kit (eBioscience). Samples were run on a BD LSRII and analyzed with FlowJo.

### Microbiota density and absolute abundances

We define microbiota density as the total DNA extracted from each sample (in *μ*g) per mg of fresh sample. For samples processed with the DIB-based extraction method, the total DNA extracted is adjusted by the fraction of the supernatant that was subsampled in the DNA extraction (e.g. a 100 *μ*L subsample is 1/7th of the total volume; total sample DNA is [DNA eluted] * 7). We then are able to utilize this measurement of microbiota density to compute the absolute abundance of microbial taxa by scaling the relative abundances of microbes in a sample by the microbiota density of that sample.

### Phylogenetic relatedness of mammalian samples

Phylogenetic relatedness was measured using sequence distance of the mitochondrial DNA sequences. All sequences were downloaded from the RefSeq organelle genome resource database (https://www.ncbi.nlm.nih.gov/genome/organelle/). Accession numbers for specific sequences used can be found in Table S1. Sequence alignment and distance measurement was performed using Clustal Omega (*55*).

### 16S rRNA data analysis

Paired end reads were joined into a single DNA sequencing using the FLASH algorithm (*56*). We split our pooled sequencing library by index using QIIME v 1.9.1 (*1*), and picked OTUs against the greengenes reference database 13_8 at 97% sequence identity (*57, 58*). The resulting OTU tables were subsequently analyzed in R (*59*) with the help of the *phyloseq* package (*60*), and custom functions developed to convert relative abundances into absolute abundances using microbiota density data.

### Statistical Analysis

Data presented were analyzed and visualized using the R statistical software (*59*). Statistical tests were used as described in the main text. For nonparametric statistical tests, multiple comparisons were performed using Dunn’s test following Kruskal-Wallis using the *FSA* R package (*61*), and corrected for multiple comparisons using Bonferonni correction. For many-to-one comparisons (e.g. pharmacologic treatments compared to untreated controls), multiple hypothesis testing correction was accomplished by using Dunnett’s test, implemented with the *multcomp* R package (*62*). For multiple comparisons between experimental groups, Tukey’s honest significant difference (HSD) was used to correct for multiple testing. Unless otherwise noted, figures depict individual samples as points, and the bars indicate the median or mean ± SEM. In figures, *p < 0.05, ** p < 0.01, and ***p < 0.001.

### Repeated sampling of gnotobiotic mice

For the experiments in which gnotobiotic mice were used to assess the roles of host carrying capacity and microbiota fitness in shaping microbiota density, mice were sampled longitudinally to increase sample size for each condition. For the mice colonized with fecal samples from the lion, elephant, ferret, and red panda, two-way ANOVA shows that the main effect is the microbiota used to colonize the mouse (F = 32.3, p = 8.27 × 10^−16^), while the identity of the individual mice does not contribute to the effects (F = 1.08, p = 0.388). The same is true for the mice colonized with fecal samples from individuals with IBD and pouch (F = 29.4, p < 0.0001 for the colonizing microbiota; F = 0.746, p = 0.634; two-way ANOVA). As a result, we are able to effectively measure the microbiota density of gnotobiotic mice in these conditions and increase the utility of each gnotobiotic mouse.

## Supplemental Results

### DNase Inactivation Buffer vs Phenol Chloroform DNA extraction comparison

To test if the two DNA extraction methods affected the resulting microbiota composition data, we processed separate aliquots from the same fecal sample using both methods. We found that the abundances of taxa in the sample processed with both methods were highly correlated (Figures S1A and S1B), suggesting that they represent equivalent ways to assay microbial community composition. In practice, the DIB method was most conducive to the small feces produced by mice and the large majority of mouse samples for this study were processed using this protocol, since the protocol utilizes smaller tubes that can be arrayed into standard 96-well formats. For the remaining mammals, the phenol:chloroform method was used as the number of stools used in the study was less, and the larger stools were more practical to aliquot into the wider 2.0 ml tube used for the phenol:chloroform method.

One possible limitation of using DNA content as a measurement of microbiota density is that small amounts of fecal matter contain sufficient DNA to saturate or clog the DNA binding columns used during extraction. This upper limit can largely be avoided by limiting the amount of input fecal material of higher microbiota density mammals (*e.g*. mice) to <50 mg and lower microbiota density mammals (*e.g*. humans) to <200 mg. In our experience, bead beating also becomes inefficient at >200 mg of fecal material. In contrast to the phenol:chloroform method, the DIB extraction protocol relies on a subsampling step that provides an additional safeguard to ensure the DNA extraction does not saturate the capacity of the Qiagen DNA-binding columns. By sampling a fraction of the lysate, we can extend the upper limit of our extraction protocol. At the extreme, using a 5 ^L subsample of the lysate can increase the dynamic range by a factor of 140, which in turn implies that we can measure microbiota density for samples containing up to 1.4 mg of DNA (140 × 10 *μ*g binding capacity of columns). On the lower end of our dynamic range, dye-based methods (Qubit Hi-sensitivity) provide an accurate detection down to 0.2 ng.

### qPCR quantification of DNA origin

While the dynamic range of the DIB extraction method described above is typically sufficient for stool samples, which contain high densities of microbial DNA compared with other environments, we further extended the method with qPCR-based quantification of the V4 region of the 16S rRNA gene. Additionally, by utilizing DNA yield per fecal sample as a measure of microbiota density, we assume host DNA is a minor contributor to the total fecal DNA yield.

To quantify the amount of bacterial and mouse DNA in our samples, we targeted the V4 region of the bacterial 16S rRNA gene (*63*) and the mouse TNF*α* gene (*64*). qPCR reactions were performed in 20 *μ*L reaction volumes with final primer concentrations of 200 nM, using KAPA SYBR FAST Master Mix (2x) ROX Low (Kapa Biosystems). The thermal cycling and imaging were performed on the ViiA 7 Real-Time PCR System (Thermo Fisher).

We quantified the amount of host vs bacterial DNA in several samples by qPCR, and evaluated the qPCR performance against spike-in controls with known combinations of mouse and bacterial DNA. We found that even amongst samples with low microbial density (e.g. samples from mice treated with vancomycin), the DNA content is largely microbial (Figure S1C). We were also able to measure the presence of microbial DNA down to concentrations near 1 pg/*μ*L (Figure S1C). This allows us to measure microbial density for samples with DNA as low as 100 pg (minimum concentration 1 pg/*μ*L in a 100 *μ*L elution volume). Coupled with the ability to subsample the lysate from our DNA extraction protocol, this allows us to measure microbiota density across 5 orders of magnitude for the phenol:chloroform method and 7 orders of magnitude with the DIB protocol.

### Absolute microbial dynamics and alpha diversity in response to pharmacologics

Culture-independent measurements have revealed that antibiotics can disrupt the composition of a healthy gut microbiota (*65*). We hypothesized that antibiotics may also have an impact on the gut microbiota density. To test this hypothesis, we administered vancomycin in two doses (0.2 mg/mL and 0.5 mg/mL) to two sets of SPF C57BL/6J mice and collected fecal pellets before and during treatment. We found that vancomycin exerted selective pressure against susceptible organisms leading to a relative expansion of Verrucomicrobia and Firmicutes in the low and high dose groups respectively (Figures S3A and S3B). When we multiplied each taxa’s relative abundance by the microbiota density to calculate their absolute abundances, we observed a bloom of Verrucomicrobia in the low dose group (Figure S3C). Surprisingly, in the high dose group, we found that vancomycin successfully depleted members of all phyla, including Firmicutes (Figure S3D). Microbiota density and alpha diversity were not significantly correlated (*ρ* = 0.107; p = 0.557; Spearman; Figure S3E), as both low dose and high dose vancomycin significantly reduced alpha diversity (p_low_ = 6.10 × 10^−5^ and p_high_ = 0. 00223, final timepoint vs baseline, Mann Whitney; Figure S3F), while only high dose vancomycin reduced microbiota density (p_low_ = 0.669, p_high_ = 0.0127, final timepoint vs baseline, Mann Whitney; Figure S3G).

## Supplemental figures

**Figure S1.**
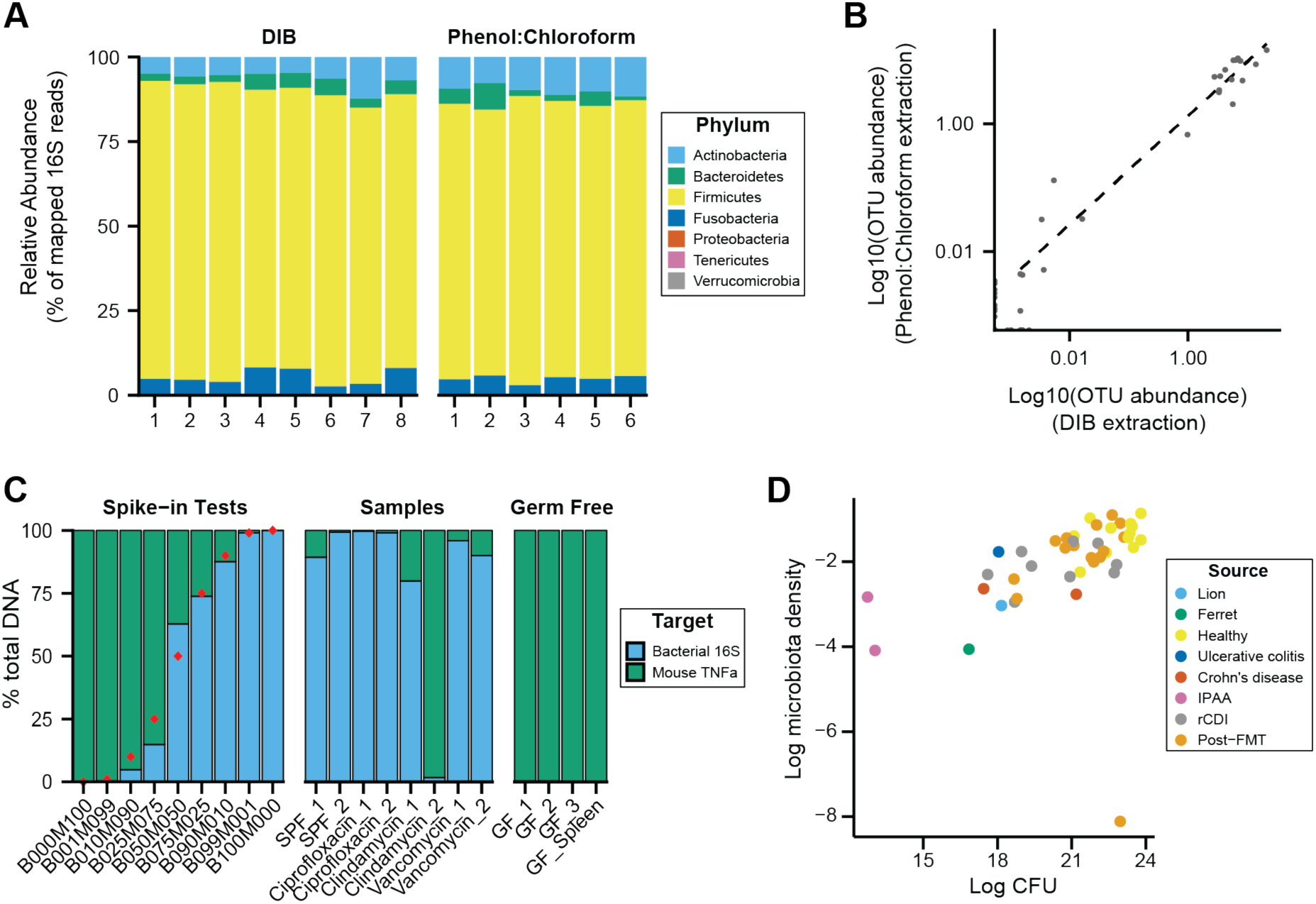
DNAse Inactivation Buffer DNA extraction method (DIB), phenol:chloroform extraction, and culture-based measurements of microbiota density yield consistent results. We homogenized one dog fecal sample and created multiple aliquots for DNA extraction using either the DIB or phenol:chloroform extraction methods. (**A**) We do not observe evidence of bias introduced by the DNA extraction method chosen, as we observe similar microbial compositions for the multiple aliquots, regardless of extraction method. (**B**) Relative OTU abundances from DIB and phenol:chloroform extracted samples are highly correlated (*ρ* = 0.904, p = 1.82 × 10^−45^, Pearson’s correlation). Dots represent average values of an individual OTU abundance across several aliquots processed using each method (n = 8 for DIB, n = 6 for phenol:chloroform). (**C**) We performed qPCR of host and bacterial fractions of mixed mouse/microbial DNA samples. Spike-in samples with known fractions of mouse and bacterial DNA (*e.g*. B010M090 = 10% bacterial + 90% mouse) were quantified with qPCR to validate the potential to identify the origin of DNA in a mixed sample. Samples from mouse fecal pellets across a variety of conditions show that the host contribution to the extracted DNA is small, even for samples with low microbiota density. Red points indicate the true spike-in percentage of bacterial DNA. GF_1, GF_2, GF_3 are host DNA controls of germ-free mouse feces. GF_Spleen is a host DNA control from a germ-free mouse spleen. (**D**) Estimates of microbiota density based on DNA content are correlated with estimates of microbiota density based CFUs from anaerobic culturing of fecal samples (*ρ* = 0.628, p = 1.05 × 10^−5^, Spearman), regardless of the source of the fecal sample. Each dot represents one sample that was quantified in parallel by colony-forming unit assay and by DNA content quantification. Colors indicate the type of sample used.

**Figure S2.**
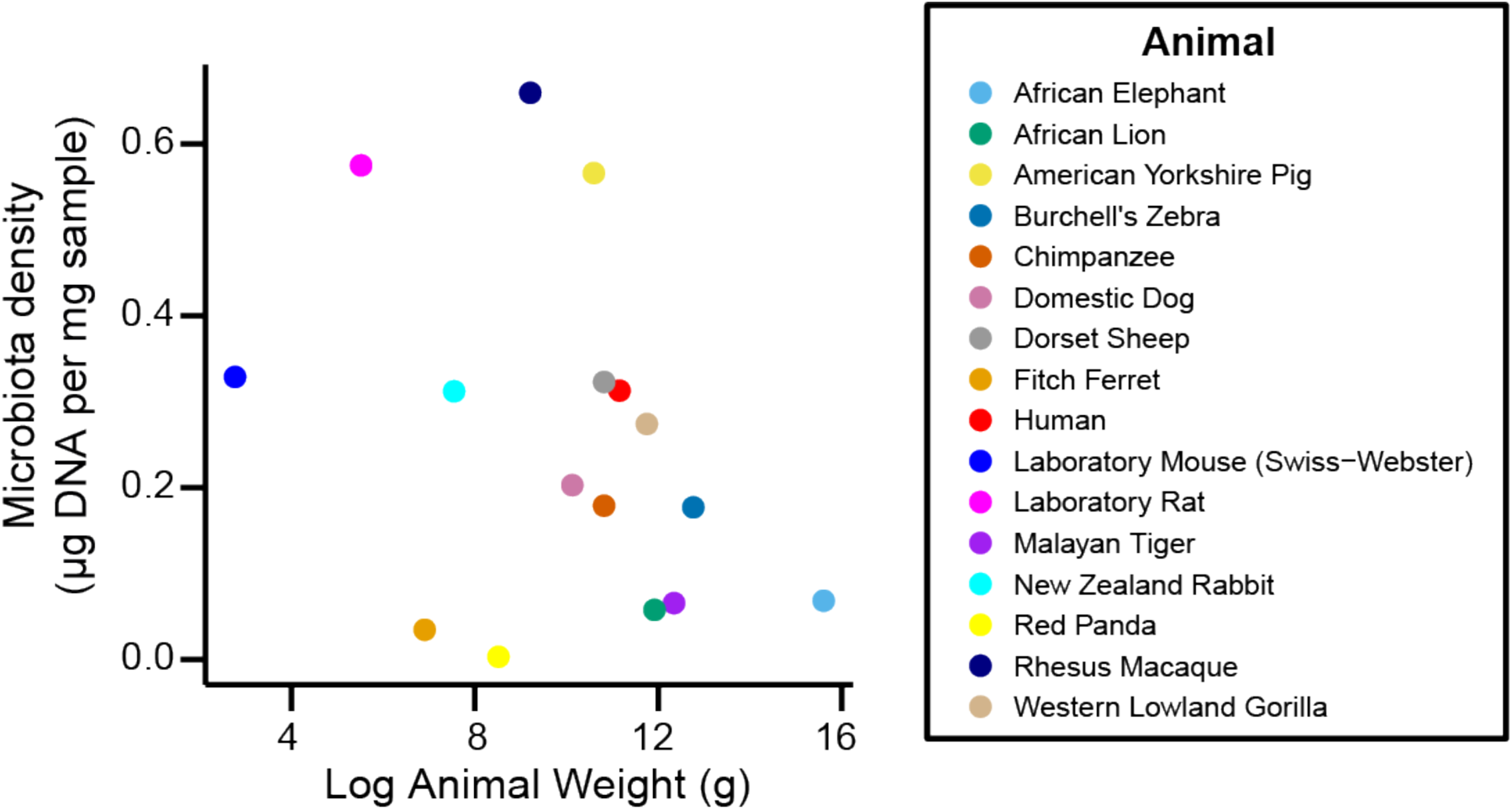
Microbiota density is not correlated with body mass.

**Figure S3.**
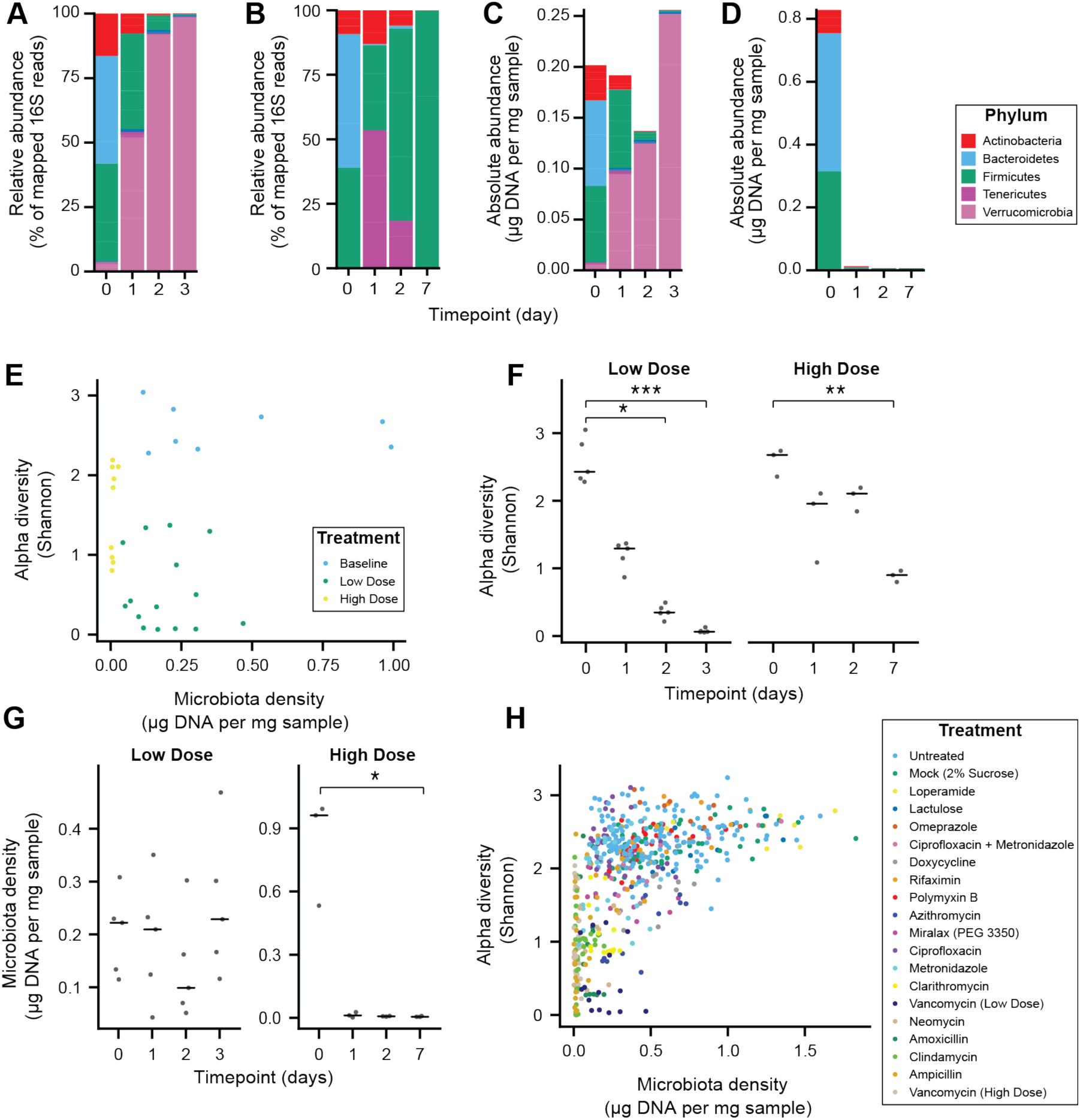
Alteration of the absolute murine fecal microbiota by pharmacologics, and the relationship between alpha diversity and microbiota density in pharmacologic interventions. (**A-D**) The relative abundances of the microbiota in SPF C57BL/6J mice treated with (**A**) low dose (n = 5) and (**B**) high-dose (n = 3) vancomycin are each dominated by a single phyla. Taking changes in microbiota density into account, the absolute abundance of the microbiota at the phylum level in the (**C**) low-dose vancomycin group demonstrates an expansion of Verrucomicrobia compared to reduction of all phyla in the (**D**) high-dose vancomycin group. (**E**) Changes in alpha diversity in response to high (0.5 mg/mL, n = 3) and low (0.2 mg/mL, n = 5) dose vancomycin treatment do not correlate with the changes observed in microbiota density. (**F**) Both low and high dose vancomycin treatment in mice reduce alpha diversity. (**G**) Low dose vancomycin did not significantly alter microbiota density, while high dose vancomycin reduced microbiota density to near zero. (**H**) Across all tested pharmacologics, there was a significant correlation between microbiota density and alpha diversity as we never observed low alpha diversity with high density. In **F** and **G**, bars indicate median, *p < 0.05, **p < 0.01, and ***p < 0.001 (Kruskal-Wallis with Dunn’s post-test corrected for multiple comparisons with the Bonferonni correction). In **E** and **H**, colors indicate treatment. For all, points represent individual samples.

**Figure S4.**
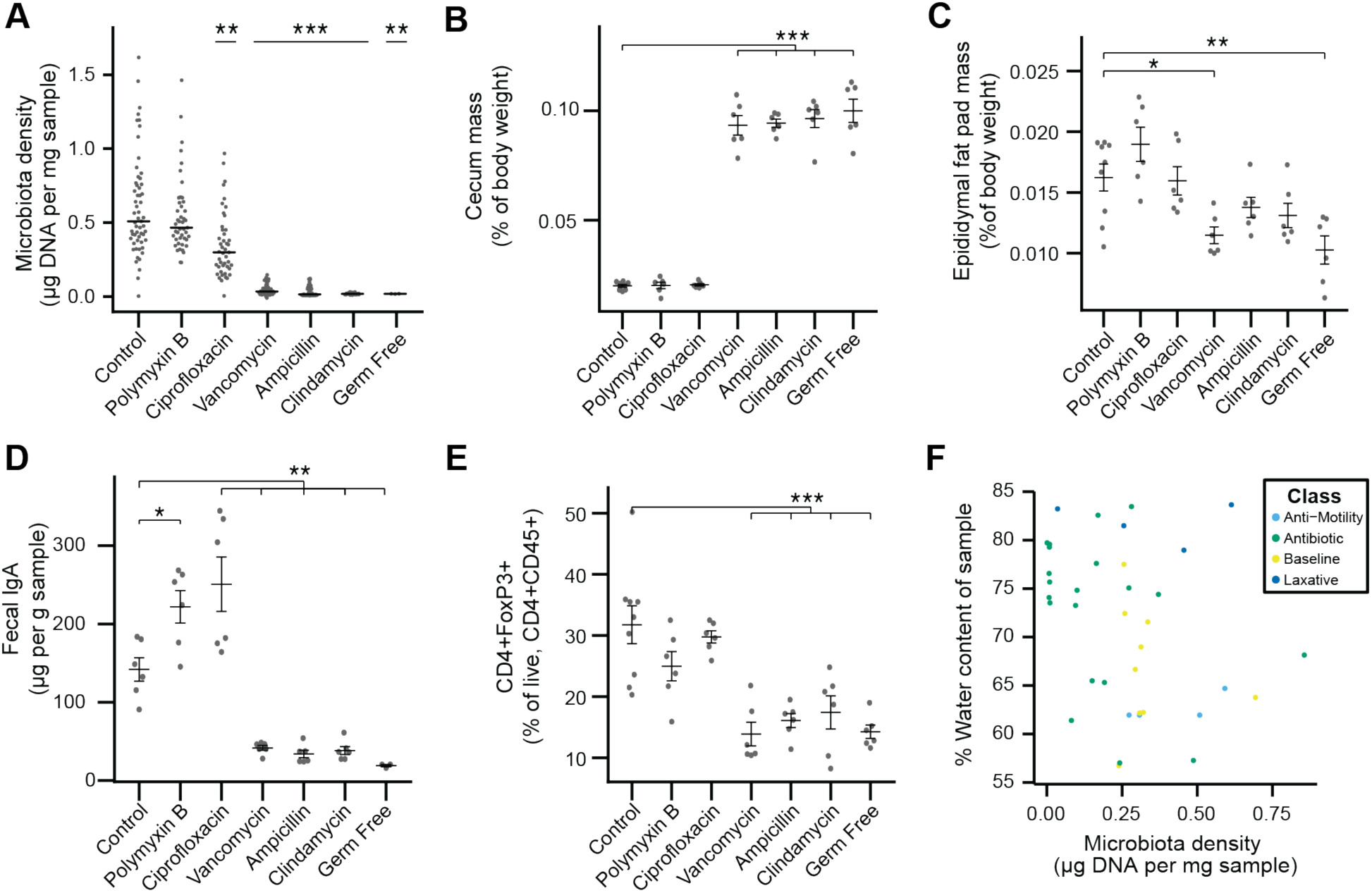
Phenotypic changes observed in antibiotic-treated mice. (**A**) Microbial density changes observed in mice administered antibiotics *ad libitum* in drinking water for four weeks. (**B-E**) The reduction in microbiota density results in changes in the (**B**) cecum size, (**C**) epididymal fat pad mass, (**D**) fecal IgA, and (**E**) colonic lamina propria FoxP3+ T regulatory cells. (**F**) Across the microbiota changes induced by the pharmacologics, microbiota density and water content are not correlated. In A, bars indicate median and nonparametric statistics used to test for significance vs control (Kruskal-Wallis with Dunn’s post-test corrected for multiple comparisons with the Bonferonni correction). In **B-E** bars indicate mean ± SEM and Dunnett’s test used to test for significance. *p < 0.05, **p < 0.01,***p < 0.001 (Dunnett’s test)

**Figure S5.**
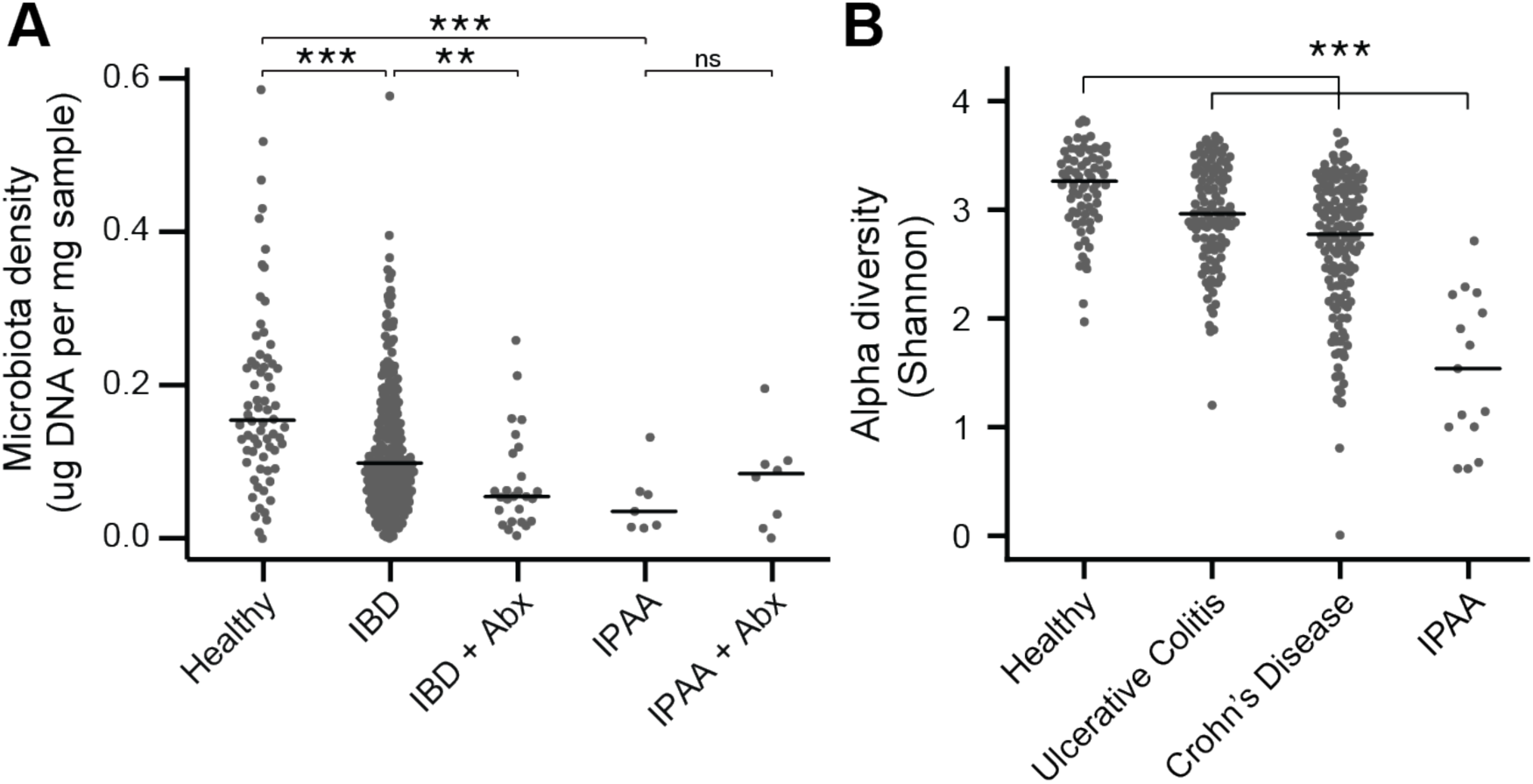
The microbiota of IBD and IPAA subjects. (**A**) Microbiota density is reduced in subjects with IBD and IPAA in the absence of antibiotic use. Nonetheless, the microbiota density of individuals with IBD on antibiotics was significantly lower for individuals with IBD on antibiotics. (**B**) Alpha diversity is reduced in subjects with IBD relative to healthy controls In **A-B**, bars indicate median, **p < 0.01, ***p < 0.001, ns = not significant.

**Figure S6.**
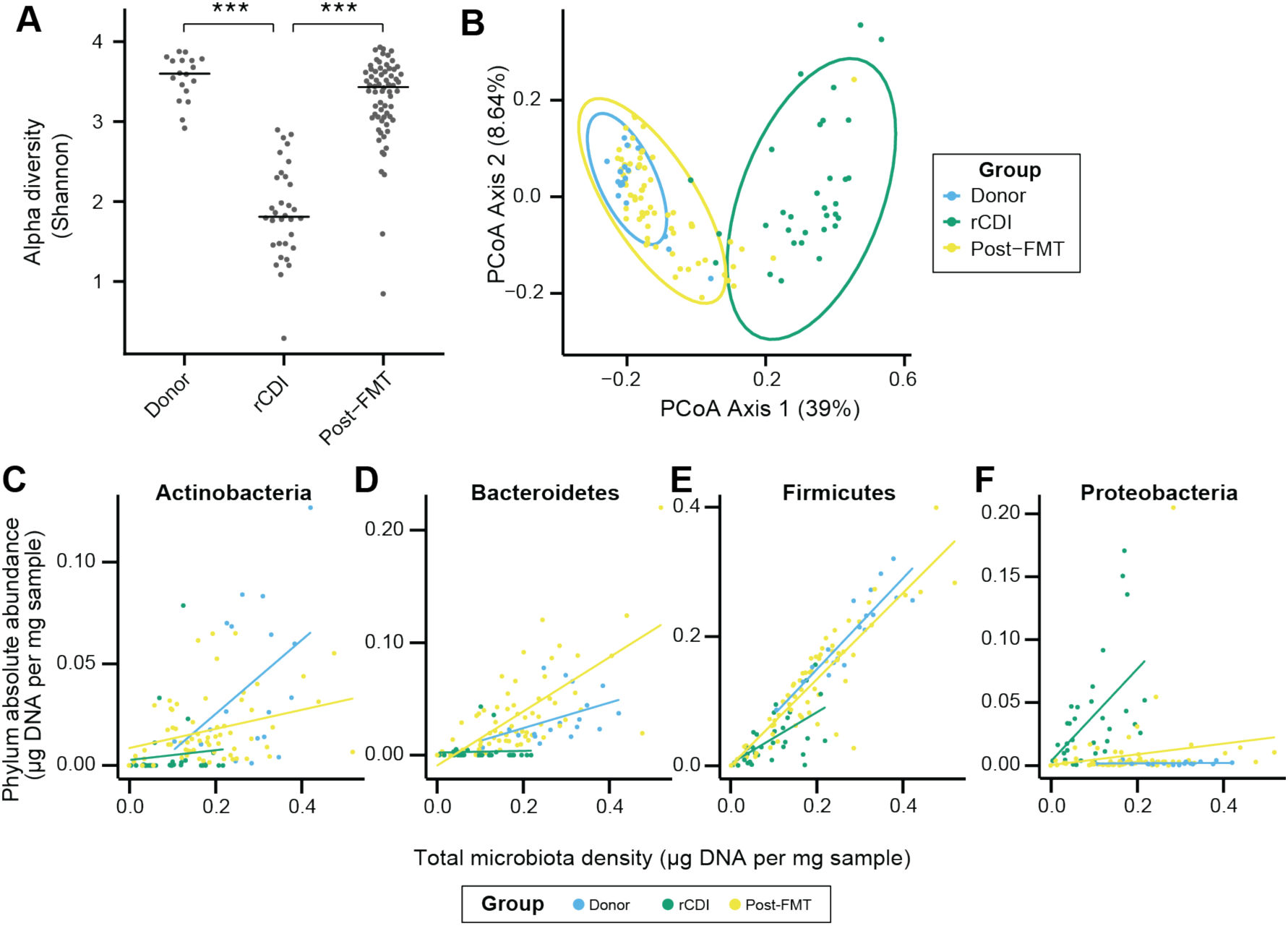
FMT changes the microbiome of individuals with rCDI to resemble that of healthy donors. (**A**) Alpha diversity in rCDI is significantly lower than in healthy individuals used as FMT donors. This change in alpha diversity is restored by FMT. (**B**) Principal coordinates analysis of unifrac distances based on the absolute abundances of OTUs in healthy FMT donors and rCDI before and after FMT. (**C-F**) The rCDI microbiota density is driven largely by the abundance of Proteobacteria and Firmicutes. In healthy donors and individuals following FMT, Proteobacteria are present at a constant absolute abundance, and microbiota density is driven by Firmicutes, Bacteroidetes, and Actinobacteria. Points represent individual subjects and colors indicate their health status In **A**, bars indicate median, ***p < 0.001. In **B**, points represent individual samples. Ellipses indicate the 95% confidence interval of distribution of points.

**Figure S7.**
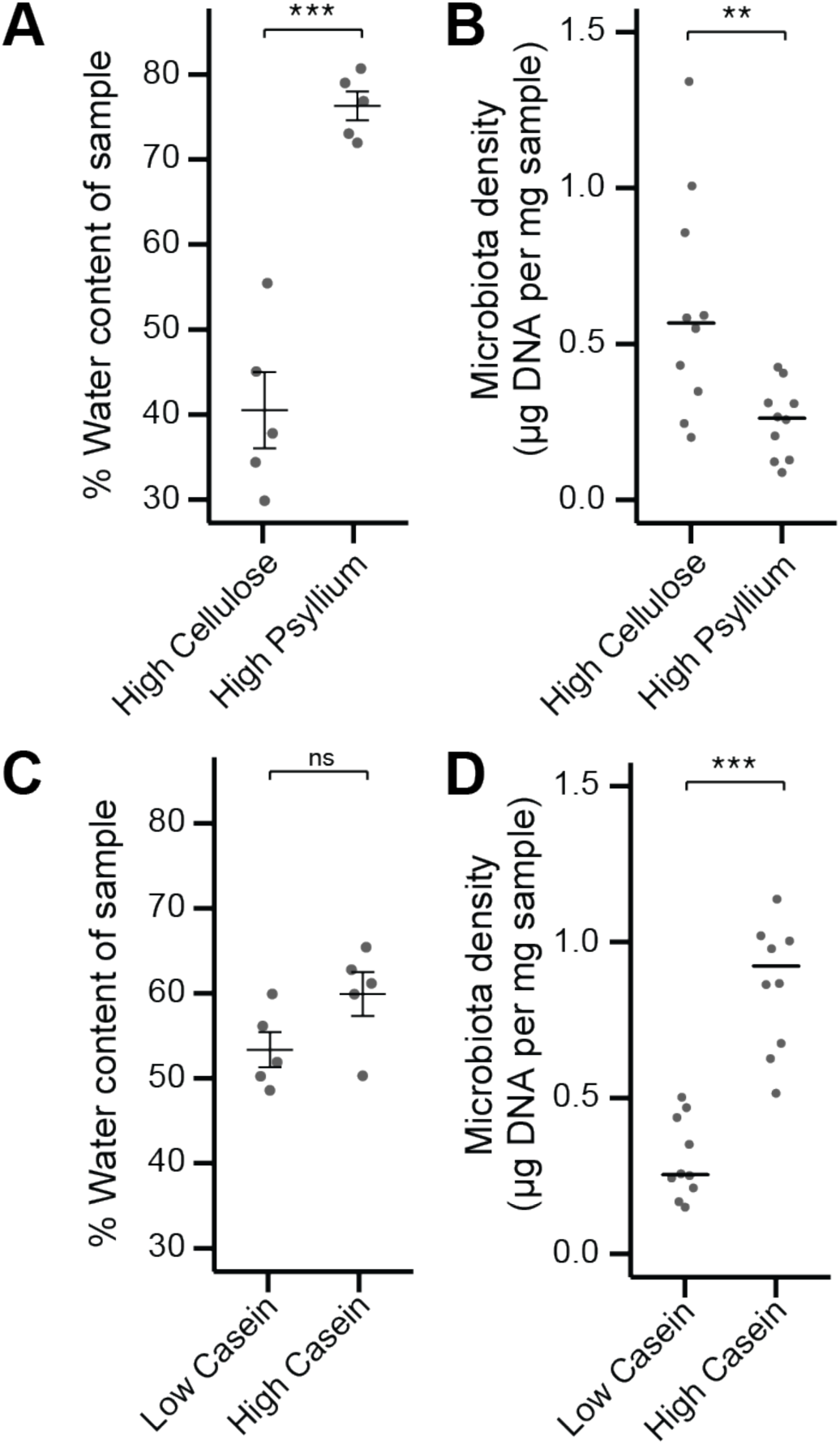
Fecal water content and microbiota density can be manipulated independently by diet. (**A**) Water content of fecal samples from mice fed diets high in soluble fiber (psyllium) is greater than that of mice fed diets high in insoluble fiber (cellulose). (**B**) There is no change in the water content of mice fed diets that vary in their protein content. (**C**) Mice fed a diet high in soluble fiber had decreased microbiota density compared to mice fed a diet high in insoluble fiber. (**D**) Protein content of the diet influences microbiota density, as shown in (*12*). In **A** and **B**, bars indicate mean ± SEM, and Student’s t test was used to test for significance. In **C** and **D**, bars indicate median, and Wilcoxon rank sum test was used to test for significance; **p < 0.01 and ***p < 0.001.

## Supplemental tables

**Table S1. Mammalian sample information.**

**Table S2. Antibiotics used in mouse experiments.**

## References

1. J. G. Caporaso et al., QIIME allows analysis of high-throughput community sequencing data. Nat. Methods. 7, 335–6 (2010).

2. P. D. Schloss et al., Introducing mothur: Open-Source, Platform-Independent, Community-Supported Software for Describing and Comparing Microbial Communities. Appl. Environ. Microbiol. 75, 7537–7541 (2009).

3. N. Segata et al., Metagenomic microbial community profiling using unique clade-specific marker genes. Nat. Methods. 9, 811–814 (2012).

4. B. M. Satinsky, S. M. Gifford, B. C. Crump, M. A. Moran, Use of Internal Standards for Quantitative Metatranscriptome and Metagenome Analysis. Methods Enzymol. 531, 237–250 (2013).

5. F. Stämmler et al., Adjusting microbiome profiles for differences in microbial load by spike-in bacteria. Microbiome. 4, 28 (2016).

6. M. A. Mahowald et al., Characterizing a model human gut microbiota composed of members of its two dominant bacterial phyla. Proc. Natl. Acad. Sci. 106, 5859–5864 (2009).

7. F. E. Rey et al., Metabolic niche of a prominent sulfate-reducing human gut bacterium. Proc. Natl. Acad. Sci. 110, 13582–13587 (2013).

8. R. Props et al., Absolute quantification of microbial taxon abundances. ISME J. 11, 584–587 (2017).

9. A. Reyes, M. Wu, N. P. McNulty, F. L. Rohwer, J. I. Gordon, Gnotobiotic mouse model of phage-bacterial host dynamics in the human gut. Proc. Natl. Acad. Sci. U. S. A. 110, 20236–41 (2013).

10. D. Vandeputte et al., Quantitative microbiome profiling links gut community variation to microbial load. Nature. 551, 507 (2017).

11. J. J. Faith, N. P. McNulty, F. E. Rey, J. I. Gordon, Predicting a human gut microbiota’s response to diet in gnotobiotic mice. Science. 333, 101–4 (2011).

12. S. R. Llewellyn et al., Interactions between diet and the intestinal microbiota alter intestinal permeability and colitis severity in mice. Gastroenterology. 0 (2017), doi:10.1053/j.gastro.2017.11.030.

13. H. Seedorf et al., Bacteria from diverse habitats colonize and compete in the mouse gut. Cell. 159, 253–66 (2014).

14. A. L. Goodman et al., Extensive personal human gut microbiota culture collections characterized and manipulated in gnotobiotic mice. Proc. Natl. Acad. Sci. U. S. A. 108, 6252–7 (2011).

15. V. K. Ridaura et al., Gut microbiota from twins discordant for obesity modulate metabolism in mice. Science. 341, 1241214 (2013).

16. G. J. Britton et al., Inflammatory bowel disease microbiotas alter gut CD4 T-cell homeostasis and drive colitis in mice. *bioRxiv*, 276774 (2018).

17. T. R. Sampson et al., Gut Microbiota Regulate Motor Deficits and Neuroinflammation in a Model of Parkinson’s Disease. Cell. 167, 1469–1480.e12 (2016).

18. G. De Palma et al., Transplantation of fecal microbiota from patients with irritable bowel syndrome alters gut function and behavior in recipient mice. Sci. Transl. Med. 9, eaaf6397 (2017).

19. E. Cekanaviciute et al., Gut bacteria from multiple sclerosis patients modulate human T cells and exacerbate symptoms in mouse models. Proc. Natl. Acad. Sci. U. S. A. 114, 10713–10718 (2017).

20. K. Atarashi et al., Induction of colonic regulatory T cells by indigenous Clostridium species. Science. 331, 337–41 (2011).

21. F. Bäckhed et al., The gut microbiota as an environmental factor that regulates fat storage. Proc. Natl. Acad. Sci. U. S. A. 101, 15718–23 (2004).

22. D. Zhang et al., Neutrophil ageing is regulated by the microbiome. Nature. 525, 528–32 (2015).

23. G. Bongers et al., Interplay of host microbiota, genetic perturbations, and inflammation promotes local development of intestinal neoplasms in mice. J. Exp. Med. 211, 457–72 (2014).

24. J. J. Faith, P. P. Ahern, V. K. Ridaura, J. Cheng, J. I. Gordon, Sci. Transl. Med., in press, doi:10.1126/scitranslmed.3008051.

25. M. B. Geuking et al., Intestinal Bacterial Colonization Induces Mutualistic Regulatory T Cell Responses. Immunity. 34, 794–806 (2011).

26. I. I. Ivanov et al., Induction of Intestinal Th17 Cells by Segmented Filamentous Bacteria. Cell. 139, 485–498 (2009).

27. A. Mortha et al., Microbiota-Dependent Crosstalk Between Macrophages and ILC3 Promotes Intestinal Homeostasis. Science (80-.). 343, 1249288–1249288 (2014).

28. P. A. Muller et al., Crosstalk between muscularis macrophages and enteric neurons regulates gastrointestinal motility. Cell. 158, 300–13 (2014).

29. Wostmann B, E. Bruckner-Kardoss, Development of cecal distention in germ-free baby rats. Am. J. Physiol. 197, 1345–6 (1959).

30. D. N. Frank et al., Molecular-phylogenetic characterization of microbial community imbalances in human inflammatory bowel diseases. Proc. Natl. Acad. Sci. 104, 13780–13785 (2007).

31. D. Gevers et al., The treatment-naive microbiome in new-onset Crohn’s disease. Cell Host Microbe. 15, 382–92 (2014).

32. U. Gophna, K. Sommerfeld, S. Gophna, W. F. Doolittle, S. J. O. Veldhuyzen Van Zanten, Differences between tissue-associated intestinal microfloras of patients with Crohn’s disease and ulcerative colitis. J. Clin. Microbiol. 44, 4136–4141 (2006).

33. J. P. Jacobs et al., A Disease-Associated Microbial and Metabolomics State in Relatives of Pediatric Inflammatory Bowel Disease Patients. Cell. Mol. Gastroenterol. Hepatol. 2, 750–766 (2016).

34. A. M. Seekatz et al., Recovery of the gut microbiome following fecal microbiota transplantation. MBio. 5, e00893–14 (2014).

35. V. Shankar et al., Species and genus level resolution analysis of gut microbiota in Clostridium difficile patients following fecal microbiota transplantation. Microbiome. 2, 13 (2014).

36. C. F. Maurice, H. J. Haiser, P. J. Turnbaugh, Xenobiotics shape the physiology and gene expression of the active human gut microbiome. Cell. 152, 39–50 (2013).

37. P. I. Costea et al., Towards standards for human fecal sample processing in metagenomic studies. Nat. Biotechnol. 35, 1069–1076 (2017).

38. R. Sinha et al., Assessment of variation in microbial community amplicon sequencing by the Microbiome Quality Control (MBQC) project consortium. Nat. Biotechnol. 35, 1077–1086 (2017).

39. G. Falony et al., Population-level analysis of gut microbiome variation. Science. 352, 560–4 (2016).

40. D. Vandeputte, G. Falony, K. D’hoe, S. Vieira-Silva, J. Raes, Water activity does not shape the microbiota in the human colon. Gut. 66, 1865–1866 (2017).

41. E. D. Sonnenburg et al., Diet-induced extinctions in the gut microbiota compound over generations. Nature. 529, 212–215 (2016).

42. M. S. Roberts, J. L. Gittleman, Ailurus fulgens. Mamm. Species, 1–8 (1984).

43. P. R. S. Blandford, Biology of the Polecat Mustela putorius: a literature review. Mamm. Rev. 17, 155–198 (2018).

44. T. Garland, The relation between maximal running speed and body mass in terrestrial mammals. J. Zool. 199, 157–170.

45. L. J. E., Primate digestion: Interactions among anatomy, physiology, and feeding ecology. Evol. Anthropol. Issues, News, Rev. 7, 8–20 (1998).

46. P. Ebinger, A cytoarchitectonic volumetric comparison of brains in wild and domestic sheep. Z. Anat. Entwicklungsgesch. 144, 267–302 (1974).

47. R. J. Smith, W. L. Jungers, Body mass in comparative primatology. J. Hum. Evol. 32, 523–559 (1997).

48. R. P. Hirten et al., Microbial Engraftment and Efficacy of Fecal Microbiota Transplant for Clostridium difficile Patients With and Without IBD. *bioRxiv*, 267492 (2018).

49. J. J. Faith et al., The long-term stability of the human gut microbiota. Science. 341, 1237439 (2013).

50. R. T. Hinnant, M. M. Kothmann, Collecting, drying, and preserving feces for chemical and microhistological analysis. J. Range Manag. 41, 168–171 (1988).

51. H. U. Bryant, C. C. Kuta, J. A. Story, G. K. W. Yim, Stress- and morphine-induced elevations of plasma and tissue cholesterol in mice: reversal by naltrexone. Biochem. Pharmacol. 37, 377780 (1988).

52. P. C. Kashyap et al., Complex interactions among diet, gastrointestinal transit, and gut microbiota in humanized mice. Gastroenterology. 144, 967–77 (2013).

53. H. Larsson et al., Inhibition of gastric acid secretion by omeprazole in the dog and rat. Gastroenterology. 85, 900–7 (1983).

54. S. Vaishnava et al., The Antibacterial Lectin RegIII Promotes the Spatial Segregation of Microbiota and Host in the Intestine. Science (80-.). 334, 255–258 (2011).

55. F. Sievers et al., Fast, scalable generation of high-quality protein multiple sequence alignments using Clustal Omega. Mol. Syst. Biol. 7, 539 (2011).

56. T. Magoč, S. L. Salzberg, FLASH: fast length adjustment of short reads to improve genome assemblies. Bioinformatics. 27, 2957–63 (2011).

57. T. Z. DeSantis et al., Greengenes, a Chimera-Checked 16S rRNA Gene Database and Workbench Compatible with ARB. Appl. Environ. Microbiol. 72, 5069–5072 (2006).

58. D. McDonald et al., An improved Greengenes taxonomy with explicit ranks for ecological and evolutionary analyses of bacteria and archaea. ISME J. 6, 610–8 (2012).

59. R Core Team, R: A language and environment for statistical computing (2017), (available at https://www.r-project.org/).

60. P. J. McMurdie, S. Holmes, phyloseq: an R package for reproducible interactive analysis and graphics of microbiome census data. PLoS One. 8, e61217 (2013).

61. D. H. Ogle, FSA: Fisheries Stock Analysis (2018).

62. T. Hothorn, F. Bretz, P. Westfall, Simultaneous Inference in General Parametric Models. Biometrical J. 50, 346–363 (2008).

63. D. A. Relman, T. M. Schmidt, R. P. MacDermott, S. Falkow, Identification of the Uncultured Bacillus of Whipple’s Disease. N. Engl. J. Med. 327, 293–301 (1992).

64. A. Nitsche et al., Quantification of human cells in NOD/SCID mice by duplex real-time polymerase-chain reaction. Haematologica. 86 (2001).

65. L. Dethlefsen, S. Huse, M. L. Sogin, D. A. Relman, The pervasive effects of an antibiotic on the human gut microbiota, as revealed by deep 16S rRNA sequencing. PLoS Biol. 6, e280 (2008).

